# Information content of cheek and lip colour in relation to the timing of ovulation in women

**DOI:** 10.1101/859801

**Authors:** Lucie Rigaill

## Abstract

Various animal species have evolved a sexual communication system with females displaying and males discriminating information about the timing of ovulation through sexual signals. More research is now investigating the potential ovulatory signalling function of female red skin colour in human and non-human primates. However, to date it is still challenging to draft satisfying hypotheses about the evolution and function of female red skin colour, due to methodological discrepancies between human and non-human primate studies. The present study used a within-individual design and objective methods to analyse the relationship between fine-scale variation in cheek and lip colour (luminance and redness) and the estimated day of ovulation in 15 cycling women. Lip, but not cheek, colour appeared to contain information about the timing of ovulation, with lips getting darker around ovulation. This study adds to the growing evidence that female red skin colour may play a role in sexual signalling in human and non-human primates but also underlines variation in trait forms and functions at the species-level.

## Introduction

Do females display visual information about their reproductive status that can modulate mate attraction and mating strategies? This question is central to sexual selection theory. The number of offspring a female can have is constrained by physiological factors such as menstrual cyclicity, gestation and post-partum amenorrhea, as well as reproductive senescence. Reproduction is usually more costly for females due to the production of larger gametes, extended maternal care, and male monopolization [1,2]. Thus, in several species, females appear to have evolved traits which are attractive to males and can act as probabilistic signals of ovulation to maximize reproduction while balancing its associated costs. Across human and non-human primates (hereafter, primates), there is evidence that female traits influence male behaviours, suggesting common evolutionary pathways and underlying mechanisms for sexual signalling [3,4].

Among the different female traits, there is a growing interest in the potential role of red skin colour in primate sexual communication. Intra-cycle variation in oestrogens induces ovulation and also affects some chromatic (redness) and achromatic (luminance) parameters of skin colour [5,6]. Circulating oestrogens bind to receptors in the skin, causing an increase in blood flow and consequently a decrease in the perceived skin luminance (i.e., darkening of the skin) [7,8]. Increase in blood flow can modulate the ratio of oxygenated/deoxygenated blood in the skin vessels which may influence perceived redness. Primate studies provide evidence that facial skin colour varies across the cycle in mandrills [9], informs about the probability of ovulation in rhesus macaques [10,11], and simultaneously conceals ovulation and advertises pregnancy in Japanese macaques [12,13]. Moreover, this colourful trait appears to be attractive to males suggesting a role in mate attraction, at least in macaques [14].

Cheek and lip colour also correlate with female attractiveness in human [15–18]. Skin colour may thus be involved into human mate attraction, although the colour of the stimuli was artificially manipulated in previous studies, which may impair the ecological validity of the results. Studies of the potential signalling function of red skin colour in women yielded mixed results. Oberzaucher et al. [19] described that cheeks were redder around ovulation compared to the end of the cycle while Samson et al [20] could not replicate these findings. Recently, Burris et al. [21] found no intra-cycle variation in cheek colour. Interpretation of these findings is constrained by some methodological limitations. While most studies used photography to analyse skin colour, which provides accurate measurements, they did not always correct lighting conditions across photography sessions using colour standards, which may have altered measurements [22,23]. Human studies usually rely on a restricted data set to assess intra-cycle difference: e.g. 2 samples from mid and late cycle, which can underestimate or overestimate cycle effects (but see [21]). Finally, human studies often failed to confirm ovulation with sex hormone profiles (again with the exception of [21]), especially when studying perceptions of intra-cycle variation. Taken together, these methodological limitations constrain our understanding of the potential role of female red skin colour in sexual signalling in humans.

To determine if women display a colourful trait that may play a role in ovulatory signalling, there is a fundamental need for studies of the relationship between fine-scale variation in trait expression and the probability of ovulation. Such studies would benefit from objective and quantitative methods, inspired by primate studies, using standardized photography, regular biological sampling, and hormonal estimation of the ovulation date [23–26]. Following this premise, this study aims at investigating whether cheek and lip colour contain information about intra-cycle variation in the probability of ovulation in women. If cheek or lip colour is related to the timing of ovulation within a cycle, I predict that cheeks or lips will be darker/redder around this timing.

## Methods

### Ethic statement

This study was approved by the Human Research Ethics Committee of the Kyoto University Primate Research Institute (KUPRI, project number 2017-13). All participants signed a written agreement concerning their participation in the study, confidentiality, and the use of their photographs for research purposes and illustrations. Participants received financial compensation in return for their participation (5,000 JPY ∼ 45 USD).

### Participants

In total, 18 women participated in the study (mean age = 28.3 ± SD 4.3 years, “Asian” = 11, “Caucasian” = 7, additional information on the supplementary material). All participants were naturally cycling and not taking hormonal contraceptives for at least 3 months prior to and during the study. I collected digital photographs and saliva samples every two days, excluding weekends and national holidays, for the duration of one complete menstrual cycle (i.e., from the beginning of their menstruations until the next ones) between September 2018 and June 2019, to limit the effect of tanning. Sampling occurred at a fixed time of the day for each participant to control for possible diurnal variation in sex hormone concentrations [27]. I collected a total of 204 photographs and saliva samples (mean par participant = 11.3 ± SD 1.5, range = 8-13).

### Assessment of cheek and lip colour

Participants removed face and lip make-up a minimum of 30 min before sampling. A single female experimenter (LR) photographed the participants, as the sex of the photographer can influence participant facial temperature and potentially skin colour [28]. Sampling was conducted in a unique location, i.e., an office room with closed curtains, ceiling lights turned on, and a constant 24° C temperature. Participants sat in front of a beige background and adopted a neutral expression. The camera was placed 2m from the participant’s chair and held at the same height as the participant’s face. The experimenter used a Nikon 5000D camera with a 12.3 megapixel CMOS sensor and a Nikon AF-S NIKKOR 18-55mm f/3.5-5.6G lens. Photographs were collected in NEF format, i.e., Nikon’s RAW format, with the flash disabled, the shutter speed and aperture size determined automatically by the camera, and a 55mm focal to limit distortion [29]. The experimenter standardised photographs by manual setting the white balance using a X-Rite White Balance Card (GretagMacbeth ColorChecker) and by photographing the participant’s face along with a Gretag X-Rite Color Checker (colour standards). This technique standardizes photographs such that colour measurements are comparable across all photographs [23–25]. Photographs were converted to uncompressed 16-bit TIFF files. Cheek colour was measured from a pair of points located on the area described in [21]. Lip colour was measured from the whole lip skin (supplementary material, Figure S1). CIELAB values were extracted using Colourworker software (Chrometics Ltd. available at: http://www.colourworker.com/) which provided luminance (L*) and red-green ratio (a*, hereafter redness) values calculated from the estimated reflectance spectrum according to the standard CIE (Commission Internationale de l’Éclairage) equations [30].

### Determination of the estimated ovulation

Participants followed the sampling guideline recommended by Salimetrics (Salimetrics, LLC, USA). Samples were collected during the photo session, labelled and stored within 15 min at - 80°C until analyses at KUPRI. Saliva samples were analysed for oestradiol (17ß-estradiol) and progesterone (4-pregenene-3,20-dione) using Salimetrics enzyme immunoassays kits. The oestradiol assay had a sensitivity of 0.1 pg/ml with an intra-assay coefficient of variation (CV) of 6.2 % and inter-assay CV of 10.8 %. The progesterone assay had a sensitivity of 5 pg/ml with an intra-assay CV of 6.6 % and inter-assay CV of 9.7 %.

The onset of the luteal phase was defined as the sample with a progesterone concentration at least 2 standard deviations greater than the mean of the 2-3 preceding baseline values [26]. Ovulation was considered to have occurred when a mid-cycle peak of oestradiol was detected around the onset of the luteal phase. Because sampling occurred every 2-3 days, I considered the day of peak oestradiol as the most likely day of ovulation and labelled it as *day 0*. The day directly preceding the estimated ovulation day was labelled as *day -1*, the day directly following it as *day* +*1*, and so on. Ovulation could not be determined (i.e., abnormal hormone variations) for 3 out of the 18 participants.

### Statistical analyses

I used photographs of 15 participants showing ovulatory cycles to analyse intra-cycle variation in cheek and lip luminance and redness (N = 165 photographs, mean per participant = 11.0 ± SD 1.4, range = 8-13).

This study tests for a possible quadratic effect of the days relative to ovulation on cheek and lip colour: higher effect toward the estimated ovulation and lower effect toward the beginning and end of the cycle. I thus constructed general linear mixed-effects models (LMMs) for cheek and lip colour (luminance and redness respectively) which included the linear (DAY) and quadratic (DAY^2^) effects of the days relative to the estimated ovulation as fixed effects, and fitted by maximum likelihood using lme4 [31] and lmerTest [32] packages in R version 3.6.0 [33]. Inspection of the cumulative distribution functions revealed good fits to the normal distribution for luminance and to the lognormal distribution for redness. Prior to modelling, the days relative to the estimated ovulation were standardized to mean = 0 and SD = 1 to improve model performance and interpretability [34]. On one hand, we can expect that the relationship between the timing of ovulation and colour would be similar across participants; I thus constructed random intercept models (RIM) with random intercepts for participant identity. On the other hand, this relationship may vary between participants; I thus constructed uncorrelated random slope models (RSM) with random slopes for the linear and quadratic terms. I also constructed the respective RIM and RSM null models in which the predictor variables were removed but the random effect structure was maintained. This resulted in 4 candidate models for each colour/skin combination: RIM-full, RIM-null, RSM-full, and RSM-null. I ensured that all relevant model assumptions were met by visually inspecting histograms of the residuals and plots of residuals against fitted values. I used an information-theory approach to objectively compare and rank the 4 candidate models in terms of how well they fitted the existing data thus assessing the likelihood that one or more models among the candidates is/are best supported by the data [35,36]. I used the function model.sel of the MuMIn package [37] to rank models based on the Akaike’s information criterion corrected for small sample size (AICc values). I reported the weight of the models which indicates to what extent one candidate model is more likely than another to provide a reasonable explanation of the variance in the data. I extracted weighted parameter estimates (β), standard errors (s.e.), and 95% confidence intervals (95% CIs) of model intercepts and predictors from conditional averaging of the 4 candidate models (function model.avg).

## Results

RIMs generally performed better than their respective RSMs (Table 1). Concerning cheek colour, the RIM-null for luminance had a weight of 0.75, while the RIM-full for redness had a weight of 0.51 (vs. 0.37 for its RIM-null), suggesting some weak evidence for variation in cheek redness only (Table 1). Concerning lip colour, the RIM-full for luminance had a weight of 0.53 (vs. 0.31 for RIM-null) and the RIM-null for redness had a weight of 0.50 (vs. 0.37 for its RIM-null), suggesting some weak evidence for variation in lip luminance only (Table 1). The timing of ovulation explained little to none of the variance in cheek redness (Table 2, Figure 1). Lip luminance varied according to the timing of ovulation with lips being darker around ovulation (ß = 0.21 ± 0.09 s.e., 95% CI = 0.03; 0.39, Table 2, Figure 2).

**Table 1.** Characteristics of candidate models for cheek and lip redness and luminance. AICc: corrected Akaike’s information criterion, weight: model probabilities

|  | logLik | AICc | weight |  | logLik | AICc | weight |
| --- | --- | --- | --- | --- | --- | --- | --- |
| *Cheek luminance* |  |  |  | *Cheek redness* |  |  |  |
| RIM - null | -247.751 | 501.7 | 0.75 | RIM - full | 279.799 | -549.2 | 0.51 |
| RIM - full | -247.292 | 505.0 | 0.14 | RIM - null | 277.351 | -548.6 | 0.37 |
| RSM - null | -247.751 | 505.9 | 0.09 | RSM - full | 279.799 | -544.9 | 0.06 |
| RSM - full | -247.292 | 509.3 | 0.02 | RSM - null | 277.621 | -544.9 | 0.06 |
| *Lip luminance* |  |  |  | *Lip redness* |  |  |  |
| RIM - full | -272.607 | 555.6 | 0.53 | RIM-null | 221.984 | -437.8 | 0.60 |
| RIM - null | -275.332 | 556.8 | 0.30 | RIM-full | 223.354 | -436.3 | 0.29 |
| RSM - null | -274.316 | 559.0 | 0.10 | RSM-null | 222.055 | -433.7 | 0.08 |
| RSM - full | -272.345 | 559.4 | 0.08 | RSM-full | 223.356 | -432.0 | 0.03 |

**Table 2.** Model averaged parameters estimates (ß) ± standard errors (s.e.) and confidence intervals (95% CIs) from conditional averaging of all candidate models. Result for which CI does not include zero are presented in bold.

|  | Intercept | DAY | DAY<sup>2</sup> |
| --- | --- | --- | --- |
| *Cheek luminance* | 26.99 ± 0.50 (**26.01; 27.97**) | -0.06 ± 0.07 (-0.21; 0.08) | 0.03 ± 0.07 (-0.11; 0.17) |
| *Cheek redness* | 2.05 ± 0.02 (**2.01; 2.10**) | 0.00 ± 0.00 (-0.00; 0.01) | 0.01 ± 0.00 (-0.00; 0.01) |
| *Lip luminance* | 20.77 ± 0.32 (**20.13; 21.40**) | 0.01 ± 0.09 (-0.18; 0.19) | 0.21 ± 0.09 (**0.03; 0.39**) |
| *Lip redness* | 2.20 ± 0.03 (**2.14; 2.27**) | 0.01 ± 0.00 (-0.00; 0.01) | 0.00 ± 0.00 (-0.00; 0.01) |

**Figure 1.**
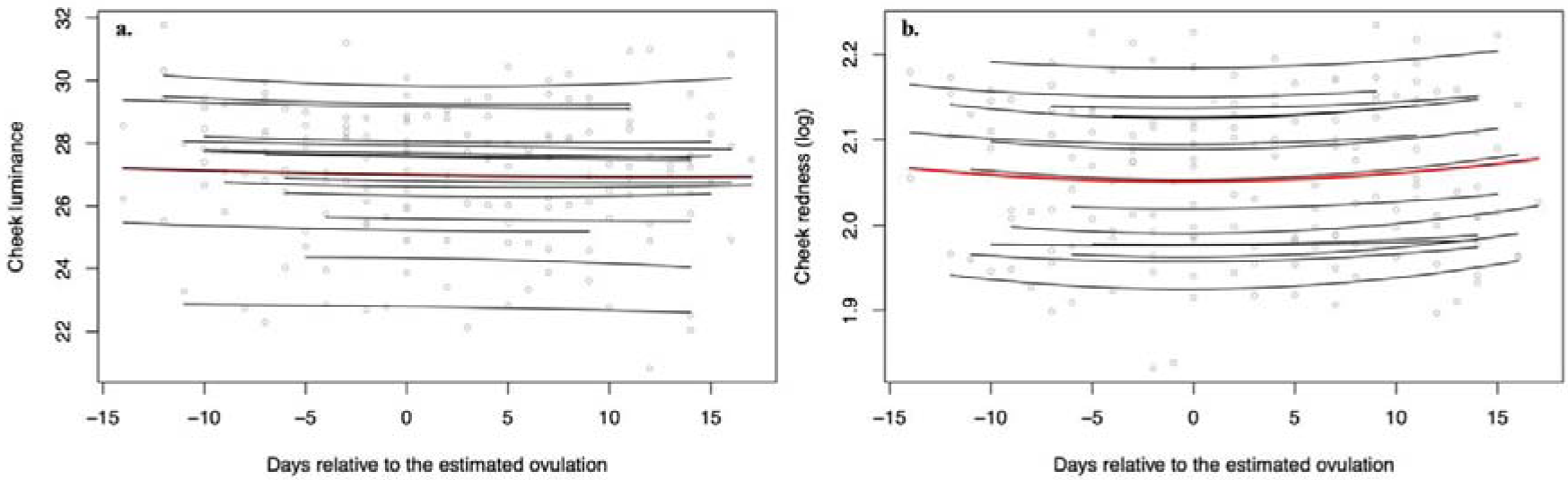
Relationship between cheek (a) luminance and (b) redness and the days relative to the estimated ovulation. Raw data are presented as grey circle. The red line presents the global model prediction; the black lines show prediction for each participant.

**Figure 2.**
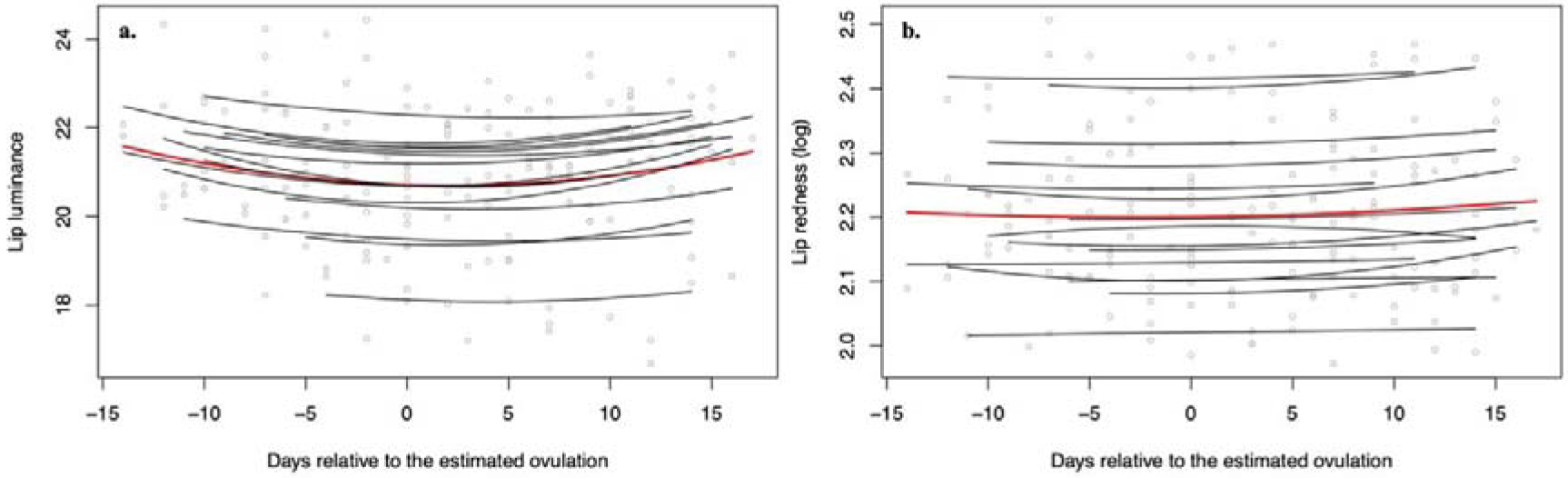
Relationship between lip (a) luminance and (b) redness and the days relative to the estimated ovulation. Raw data are presented as grey circle. The red line presents the global model prediction; the black lines show prediction for each participant.

## Discussion

Using objective methods to analyse fine-scale and intra-individual changes in cheek and lip colour according to the timing of ovulation, I found that lip luminance contains information about the timing of ovulation in women. Cheek colour may not be related to the timing of ovulation in agreement with a previous study [21].

Different facial features may respond differently to variation in the probability of ovulation. Intra-cycle variation in colour may be less cryptic for the lip skin as a result of its higher vascularization and finer cellular layers [38] or due to tissue-specific differences in oestrogen receptor expression or sensitivity. The latter has already been suggested to explain the differential effect of cycle phase on facial and hindquarter colour in Japanese macaques [13]. Further studies on tissue-specific expression and sensitivity of oestrogen receptors would help to clarify this question. It is also possible that hormonal contraceptives have on-going and long-term effect on red skin colour (e.g., female odours [39,40]), a question that I could not address in the present study and requires further investigation. Alternatively, colour changes in the facial skin may be more condition-dependent and related to individual health, thus masking a potential cycle effect [15,16,41].

So, can lip colour act as an ovulatory signal? The present study suggests that lip colour contains information about the timing of ovulation. To determine whether this information is conveyed, further studies should assess men and women responses toward intra-cycle variation in lip colour to determine whether these changes are perceptible. However, caution should be used when designing such experiments. Studies should use an intra-individual design, i.e., using individual pictures rather than pictures from different participants or composite stimuli, as the differences in skin colour appeared to be relatively greater between than within participant in the present study. Studies should also collect and test multiple samples per individual in order to detect fine-scale variation, and confirm ovulation using hormonal data or ovulation detection kit. Such experimental design should reduce the risk of over- or under-estimating a potential cycle effect. However, whether variation in this trait is detectable under “optimal” conditions, i.e., artificial environments and controlled settings, does not necessarily entail consequences on human sexual communication and mate preferences since laboratory studies can have low ecological validity. At best, studies on perception could provide further evidence that lip colour has the potential to play a role in ovulatory signalling.

Humans may have inherited the biological bases for female red skin colour and sexual signalling from a primate ancestor. Female red skin colour has been suggested to play a role in the sexual communication of some primate species, i.e., signalling [9–11] or concealing ovulation [13], and advertising pregnancy [9,12]. The differences in red skin traits (face, lips, and sexual skin) and signalling functions probably result from the different socio-environmental constraints on mating across species. Concealing the reproductive status may have evolved in species facing higher male monopolization or pair-bonding, lower intra-sexual competition, and higher costs on signalling; while higher infanticide risks and intra-sexual competition, limited mating opportunities, and lower costs on signalling may have favoured exaggerated or multimodal signalling [42,43]. However, these proposed effects do not appear to fully explain the inter-species differences observed in female red skin colour functions, as species with similar socio-ecology express traits that may be involved in either ovulatory signalling or concealing (e.g. [11,13]). More studies using longitudinal and ecologically valid designs are needed to better understand the possible evolutionary pathways and underlying mechanisms leading to the evolution of female red skin colour in primate species including humans.

## Acknowledgments

I acknowledge the Human Research Ethics Committee of Kyoto University Primate Research Institute for permission to conduct my research. I would like to sincerely thank C. Neumann for his insightful comments on the study and his valuable advice with statistical analyses, J. Duboscq, C. Garcia, and AJJ MacIntosh for comments on an earlier version of the manuscript, and K. Mouri for her general support in the lab.

This is a pre-print of an article published in Behavioral Ecology and Sociobiology. The final authenticated version is available online at: https://doi.org/10.1007/s00265-020-02851-y

